# Combined quetiapine and radiation therapy approach to treat mesothelioma-initiating cells and increase survival in a mouse model of mesothelioma

**DOI:** 10.1101/2025.04.08.647821

**Authors:** Anjelica Cardenas, Evelyn Arambula, Linda Azizi, Tara Lam, Sabrina Sutlief, Kruttika Bhat, Mohammad Saki, Ling He, Frank Pajonk

## Abstract

**Introduction:** Malignant pleural mesothelioma (MPM) is a rare thoracic cancer associated with poor prognosis and low survival rates. In solid cancers, repurposed dopamine receptor antagonists have been shown to have anti-cancer effects. Moreover, in combination with radiotherapy, quetiapine (QTP), a dopamine (D) 2/3 receptor antagonist, has been shown to interfere with self-renewal capacity in glioma-initiating cells and increase survival in mouse models of glioblastoma. In this study we explore combined treatment effects in MPM.

**Methods:** Using mesothelioma cell lines, MSTO-211H, H2052, and H2452, and a MSTO-211H-derived orthotopic xenograft mouse model of MPM we examined how QTP combined with radiation affects mesothelioma-initiating cells (MICs) *in vitro* and survival *in vivo*. Subsequently, bulk and single cell RNA sequencing was used to characterize the transcriptomic landscape of MSTO-211H treated with combined radiation and QTP.

**Results:** We demonstrate that combining QTP with radiation reduces MIC self-renewal capacity and stem cell frequency. *In vivo*, this combination therapy significantly extends the median survival of mesothelioma-bearing mice. Clonogenic survival assays revealed that QTP does not enhance radiosensitivity in the tested mesothelioma cell lines. Sequencing data revealed, combined treatment downregulated cell cycle and proliferation pathways, depleted cancer stem cells, and increased cellular senescence.

**Conclusion:** Taken together, our study highlights the therapeutic potential of radiation with QTP in the treatment of MPM.

## INTRODUCTION

Malignant pleural mesothelioma (MPM) is an aggressive form of cancer in the pleura, associated with long-term exposure to asbestos. Symptoms typically do not arise until the late stage of the disease, leading to a grim prognosis. While incidences of MPM have decreased in the United States, incidences are still prominent in developing countries ^1^. The standard treatment for MPM is surgery and platinum-based chemotherapy, however, the 5-year survival rate is approximately 12% ^2^. With MPM relatively resistant to current treatments and few alternative or new therapeutics emerging, it is critical to find novel therapeutic strategies to treat MPM.

Mesothelioma-initiating cells (MICs) have previously been characterized via sphere formation, tumor-initiating capacity, and expression of cancer stem cell markers ^3–7^. Repurposed dopamine receptor antagonists have demonstrated anticancer effects in brain, breast, colon, gastric hepatocellular, and sarcoma cancers ^8–13^. Quetiapine (QTP), a second-generation dopamine receptor antagonist, has previously shown to be efficacious in Glioblastoma (GB) in combination with radiotherapy without affecting radiation sensitivity of glioma-initiating cells (GICs) ^14^. Recently, our lab has shown dopamine receptor antagonist treatment in combination with radiation interferes with self-renewal capacity in GICs, prevents conversion of non-GICs into induced GICs, and prolongs survival in mouse models of GB ^14–16^.

Radiotherapy (RT) is not a standard treatment regimen for MPM, instead, it is typically reserved for palliation or as an adjunct to resection ^17, 18^. Unfortunately, the use of RT in treating MPM is limited by a lack of robust evidence, few studies, and minimal clinical guidelines supporting the procedure ^18, 19^. However, radiosensitivity has been observed *in vitro* ^20^. With few studies evaluating the effects of RT alone or in combination with chemotherapy in MPM, more research is necessary.

In the present study, we test whether QTP combined with radiation will impact MICs and overall survival in a mouse model of MPM. To investigate this, we examined the effects of QTP on mesothelioma cell lines using sphere formation assays, limiting dilution assays, and clonogenic survival assays. We subsequently assessed combination treatment of radiation and QTP on overall survival *in vivo*. Further, we characterized cellular responses to combined treatment using both bulk and single-cell RNA sequencing.

## MATERIALS AND METHODS

### Cell Culture

Mesothelioma cell lines MSTO-211H (biphasic histotype), NCI-H2052 (epithelial histotype), and NCI-H2452 (epithelial histotype) were purchased from American Type Culture Collection (ATCC, Manassas, VA, USA). Cell lines were cultured in log-growth phase in Roswell Park Memorial Institute (RPMI) 1640 medium (Gibco, Thermo Fisher Scientific, USA) supplemented with 10% fetal bovine serum (FBS) and 1% penicillin-streptomycin (10,000 U/mL, Gibco, Thermo Fisher Scientific, Waltham, MA, USA). Cells were propagated into MICs in non-treated plates with serum-free conditions (RPMI 1640 medium supplemented with SM1, epidermal growth factor (EGF), fibroblast growth factor 2 (bFGF), heparin, and 1% penicillin-streptomycin). Mesothelioma cells were grown in a humidified atmosphere at 37°C with 5% CO_2_. Cell line identities were confirmed by DNA fingerprinting (Laragen, Culver City, CA, USA) and routinely tested for mycoplasma infection with a mycoplasma detection PCR kit (Applied Biological Materials, Richmond, BC, Canada) and/or MycoAlert (Lonza, Walkersville, MD, USA).

### Quantitative Real-time Polymerase Chain Reaction (qRT-PCR)

Total RNA was isolated using TRIZOL reagent (Invitrogen, Thermo Fisher Scientific). SuperScript Reverse Transcription III (Invitrogen, Thermo Fisher Scientific) was used to synthesize complementary DNA. Quantitative PCR was performed in the QuantStudio^TM^ 3 Real-Time PCR System (Applied Biosystems, Carlsbad, CA, USA) using the PowerUP^TM^ SYBR^TM^ Green Master Mix (Applied Biosystems, Carlsbad, CA, USA). Ct for each gene was determined after normalization to PPIA (housekeeping gene) and Λ1Λ1Ct was calculated relative to the reference sample. Gene expression values were set equal to 2-Λ1Λ1Ct as described by the manufacturer (Applied Biosystems, Carlsbad, CA, USA). All PCR primers (**Supplementary Table 1**) were synthesized by Invitrogen.

### Limiting Dilution Assays

Cells were seeded under serum-free conditions at clonal densities into non-treated tissue culture plates. The next day, cells were treated and irradiated 1h later using an X-ray irradiator. Cells were grown for 7-14 days and supplemented with 10 µl/well of media with growth factors every other day. Upon formation of visibly distinct spheres, spheres were counted in each well using a conventional microscope and recorded. For sphere formation capacity (SFC), sphere formation at each dose was normalized against the non-irradiated control. Data points were fitted using a linear quadratic model. Furthermore, limiting dilution assays were calculated and plotted using Extreme Limiting Dilution Analysis software ^21^.

### Animals and Malignant Pleural Mesothelioma Tumor Model

Six-to eight-week-old NOD-*scid* IL2Rgamma^null^ (NSG) mice used in this study were from The Jackson Laboratories, derived, bred, and maintained in a pathogen-free Department of Radiation Oncology facility at UCLA. Mice were housed in an American Association of Laboratory Animal Care-accredited Animal facility in accordance with local and national guidelines for the care of animals. Daily weights of animals were recorded. 1 x 10^5^ MSTO-211H GFP-Luc cells in 200 µL PBS were injected into the thoracic cavity for pleural dissemination as described by Yasui and colleagues ^22^. Eight days after pleural injection, successful grafting of tumors was confirmed by bioluminescence imaging (BLI), followed by the administration of respective drug treatment and irradiation. Mice that lost >20% of body weight or difficulty breathing were euthanized.

### In Vivo Bioluminescent Imaging

On day 8 after implantation of xenografts, MSTO-211H GFP-Luc-bearing NSG mice were imaged. Tumor-associated bioluminescent signal was recorded. Mice were injected intraperitoneally (i.p.) with 100 µL of D-luciferin (15 mg/mL; Syd Labs, Hopkinton, MA prior to imaging. Ten minutes later, mice were anesthetized (2% isofluorane gas in O_2_), and luminescence was recorded (IVIS Lumina II Systems, PerkinElmer, Waltham, MA). Images were analyzed with Living Image Software (Caliper life Sciences, Hopkinton, MA).

### Drug Treatment

Following tumor grafting confirmation via BLI, mice bearing MSTO-211H tumors were injected on a 5-d on/2-d off schedule with saline or quetiapine fumarate (QTP, Key Organics, Cornwall, UK, 30 mg/kg) subcutaneously or until they reached euthanasia endpoints. QTP was dissolved in sterile saline at a concentration of 5 mg/mL.

### Irradiation

Prior to irradiation, mice were given an IP injection with a 4:1 mixture of ketamine (100 mg/mL; Phoenix, MO) and xylazine (20 mg/mL; AnaSed, IL) to anesthetize animals and placed on their left side into an irradiation jig. Mice received thoracic irradiation with lead shielding used to protect non-thoracic regions. Irradiation was delivered using an experimental X-ray irradiator (Gulmay Medical Inc.) at a dose rate of 5.519 Gy/min at room temperature. The X-ray beam was operated at 300 kV and hardened using a 4-mm Be, a 3-mm Al, and a 1.5-mm Cu filter and calibrated using National Institute of Standards and Technology-traceable dosimetry. Animals implanted with MSTO-211H cells received a single dose of 10 Gy on day 8 after tumor implantation. Corresponding controls were sham irradiated.

### Bulk RNA Sequencing

100,000 MSTO-211H MICs were seeded in suspension. The next day, cells were treated with 10 μM of QTP or vehicle control and irradiated (0 or 4 Gy). 48h post-irradiation, total RNA was isolated from cells with Trizol. Bulk RNA sequencing (Bulk RNA-seq) was performed by Novogene (Sacramento, CA) and reads were mapped to the human genome (hg38) following their standard pipeline. Raw data counts obtained from each treatment group: 0Gy_DMSO (Con), 4Gy_DMSO (IR), Gy_10μMQTP (Rx), and 4Gy_10μMQTP (Combo) were input directly into the integrated Differential Expression and Pathway analysis (iDEP2.0) webtool ^23^, for performing differential gene expression and pathway analyses. Data was originally pre-processed using the following parameters: minimum counts per million (CPM) of 0.1, in 1 library. DESeq2 was used for identifying differentially expressed genes (DEGs) with the following parameters: false discovery rate (FDR) cutoff of 0.2 and minimum fold change of 1.2. Pathway enrichment analysis was done using the hallmarks.MSigDB dataset. Downregulated genes identified were input into StemChecker ^24^. Significance (p-value) was calculated by the hypergeometric test and adjusted p-value was calculated with Bonferroni correction.

### Single Cell RNA Sequencing

MSTO-211H MICs were seeded in a flask. 24h later, cells were treated with 10 μM of QTP or vehicle control and irradiated (0 or 4 Gy) 1hr after. 48h post-irradiation, cells were prepared for single cell RNA sequencing (scRNASeq) according to PIPseq V T2 3’ Single Cell RNA Kit User Guide (Fluent Biosciences). cDNA and library quality were assessed by ThermoFisher Qubit 4 Fluorometer and Agilent 4200 TapeStation System. cDNA libraries with 20,000 cells/sample were sequenced on the Illumina 6000 system and analyzed. Clustering and cell-type annotation analysis was run with the Seurat package (v5.1) in RStudio (v4.4.2). Matrices were filtered for cells with high mitochondrial and ribosomal gene count. Briefly, highly variable genes were generated to perform principal component analysis (PCA). Groups were projected onto Uniform Approximation and Projection (UMAP) analysis run. Cluster-specific markers genes were identified via FindAllMarkers to normalized gene expression data. Cell-type identities were annotated using SingleR with HumanPrimaryCellAtlasData as a reference. SenePy ^25^ was used to evaulate cellular senescence in our scRNAseq dataset.

### Statistical Analysis

Statistical analyses were performed using the GraphPad Prism Software (GraphPad Software, San Diego, CA), ELDA webtool software ^21^, and StemChecker ^24^, and SenePy ^25^. Unless specified, data was represented as mean +/-standard error mean (SEM) of at least 3 biological replicates.

## RESULTS

### Combined QTP and radiation treatment reduces self-renewal capacity and stem cell frequency in mesothelioma

Since QTP is a dopamine (D)2/3 receptor antagonist, we first determined whether MICs express D2/3 receptors. To do this, we assessed gene expression in MSTO-211H MICs and adherent cells. We observed minimal differential gene expression in MICs versus adherent cells in the expression of DRD2 and DRD3 (**Figure 1A**). Baseline expression of additional QTP target receptors were investigated and we observed Histamine 1 receptor (HRH1) and two serotonin receptors (HTR1B and HTR1D, **Supplementary Figure 1**) were expressed in MSTO-211H MICs. Taken together, expression of target receptors suggest that MICs may be susceptible to QTP.

**Figure 1.**
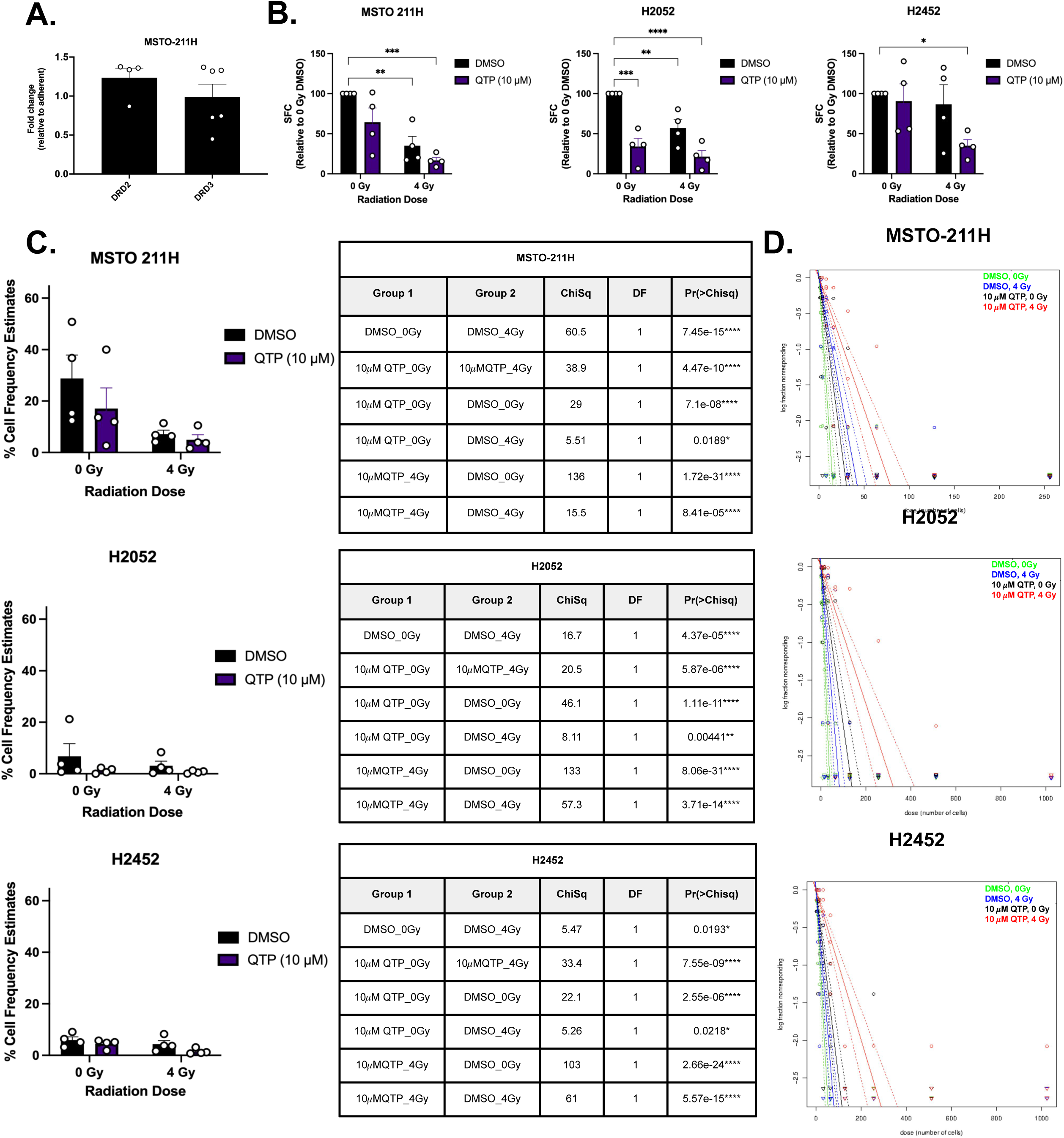
*Combined QTP and radiation treatment reduces self-renewal capacity and stem cell frequency in mesothelioma.* **A.)** Baseline expression of dopamine receptors (DRD) 2 and 3 in mesothelioma-initiating cells (MICs) relative to adherent mesothelioma cell lines (MSTO-211H, H2052, and H2452), n=4-6 biologically independent repeats. **B.)** Sphere forming capacity (SFC) in presence of QTP (10μM) and radiation (0 or 4 Gy), n=4 biologically independent repeats/ group. **C.)** Left: Bar graph of % Cell Frequency Estimates and Right: Tables of pairwise tests for differences in stem cell frequencies. **D.)** Log-fraction plots of the limiting dilution model fitted to the data. The slope of the line is the log-active cell fraction. The dotted lines give the 95% confidence interval, n= 4 biological repeats/group. All data represented as mean +/-SEM, except in the case of log-fraction plot.

Using limiting dilution assays, we then tested the effects of QTP alone and in combination with radiation (4 Gy) on self-renewal capacity and stem cell frequency *in vitro*. In three cell lines tested (MSTO-211H, H2052, and H2452), a significant decrease in SFC in combination treated groups relative to controls was observed (**Figure 1B, Supplementary Table 2**). Consistent with a decrease in self-renewal capacity, combined QTP and radiation treatment significantly decreased stem cell frequencies in all cell lines tested (**Figure 1C-D**). **Supplementary Tables 3-4** show confidence intervals for 1/(stem cell frequency and overall test for differences in stem cell frequencies between any of the groups calculated via ELDA webtool ^21^. Our findings suggest combined QTP with radiation treatment prevent MIC self-renewal capacity and reduce stem cell maintenance. To assess the effects of QTP alone and in combination with varying doses of radiation on MPM colony formation, we used clonogenic survival assays (**Figure 2A-B**). The surviving fractions (SF) were calculated by normalizing the plating efficiency of treated cells to that of corresponding control cells (0 Gy, DMSO). Compared to controls, no reduction in SF following irradiation at 2, 4, 6, and 8 Gy for three cell lines tested were observed (**Figure 2B**). Therefore, our results suggest QTP does not enhance the response to irradiation as indicated by clonogenic cell survival.

**Figure 2.**
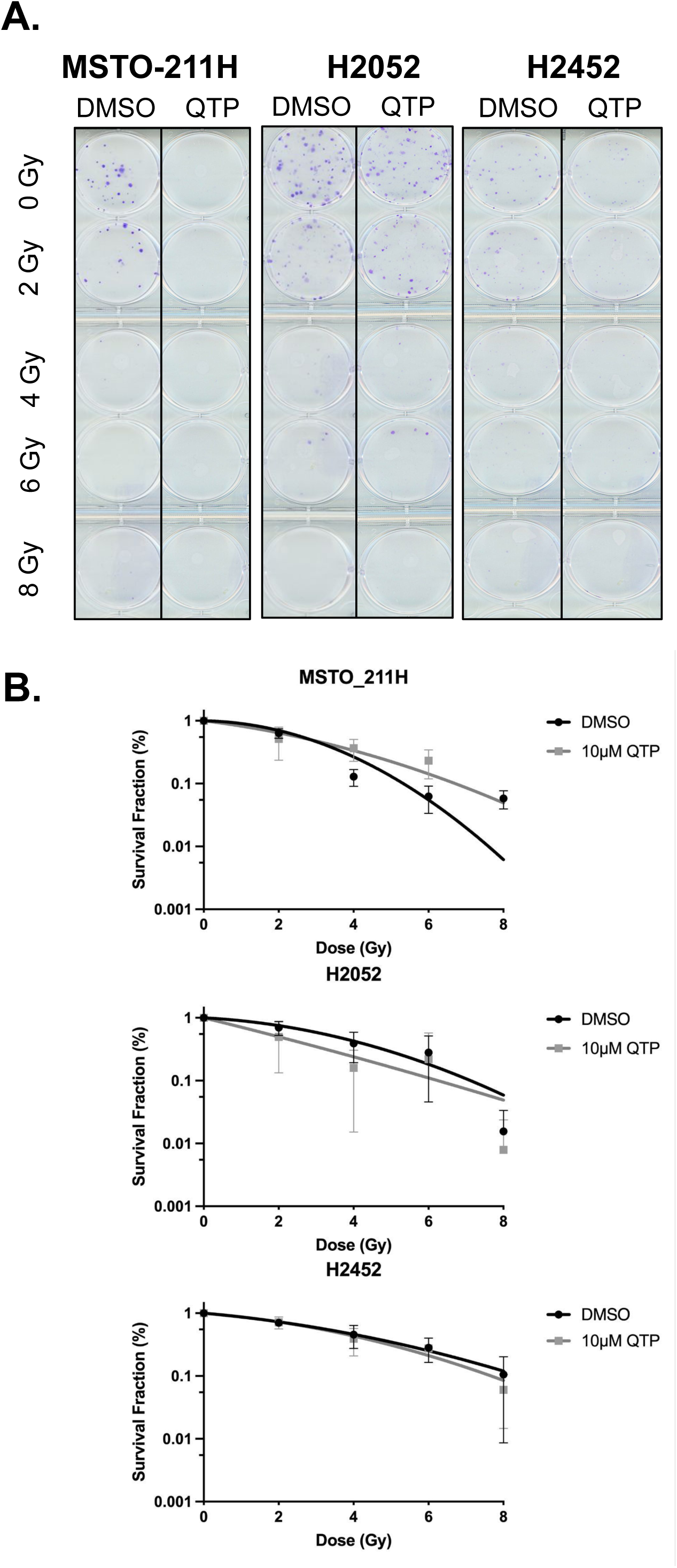
*QTP does not significantly reduced clonogenic survival after radiation.* **A.)** Representative images of colonies for all treatments and cell lines tested via clonogenic survival assays. **B.)** The surviving fractions (SF) were calculated after normalizing to the plating efficiency of the corresponding control cells. The results are represented as the mean % SF +/-SEM, n=3 biologically independent replicates/group.

### Combined QTP and radiation treatment prolongs survival in vivo

To test the combination effects of radiation and QTP on MICs *in vivo*, we implanted NSG mice with MSTO-211H into the thoracic cavity ^22^ (**Figure 3A**). After engraftment was confirmed on day 8 via BLI, mice were treated 5d on/ 2d off until they reached euthanasia criteria (**Figure 3A-B**). Our results show combined radiation and QTP treatment significantly improved the median survival compared to all groups tested (**Figure 3C, Supplementary Table 5**). Both radiation and QTP alone did not significantly improve survival. Weights were not impacted by daily QTP treatment or combined treatment (**Figure 3D**) and following euthanasia, MPM was visually confirmed (**Figure 3E**). Taken together, combined radiation and QTP treatment is effective in our orthotopic xenograft mouse model of MPM.

**Figure 3.**
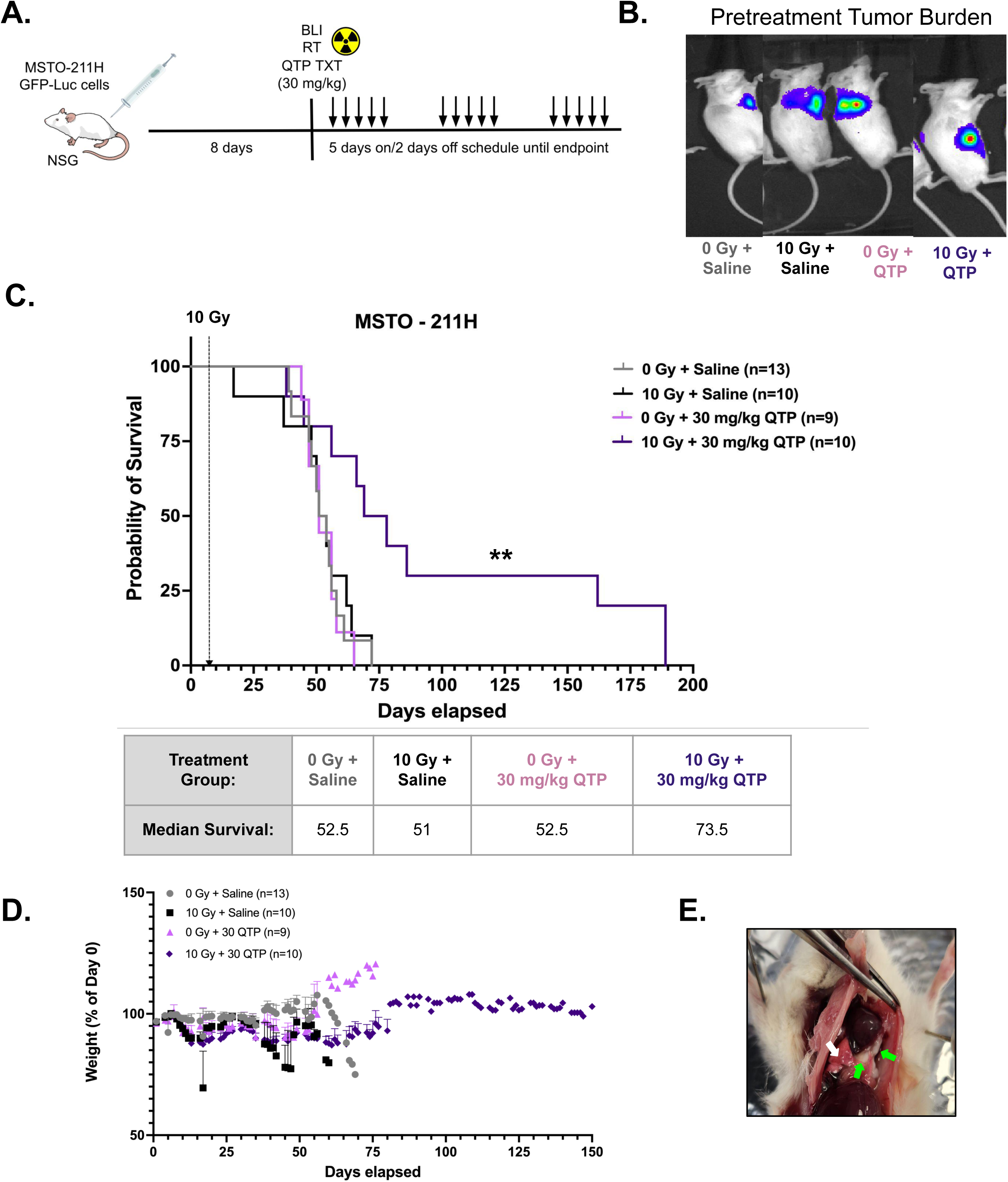
*Combined QTP and radiation treatment improves overall survival in a mouse model of MPM.* **A.)** Schematic of the treatment paradigm. **B.)** Bioluminescent imaging prior to treatment to confirm graftment on day 8. **C.)** Top: MSTO-211H tumor-bearing NSG mice were treated with a single dose of 10 Gy and/or QTP (30 mg/kg), n=9-13 mice/group. Bottom: Table of mean survival for each treatment group. **D.)** Weight graph of treated mice over time **E.)** Representative images of tumor burden in mice. White arrow indicates lung, green arrows indicate MPM mass.

### Bulk RNA Sequencing

To determine which pathways were activated in cells that were treated with combined QTP and radiation, we performed bulk RNA sequencing (bulkRNAseq) on MSTO-211H MICs. We completed sequencing in MSTO-211H because it is a mixed tumor with both epithelioid and sarcamatoid mesothelioma cell types. Differential gene expression analysis identified 298 and 395 genes upregulated by 4 Gy + QTP (Combo) relative to unirradiated, DMSO control and ionized radiation (IR, 4Gy) alone samples. 436 and 912 genes were downregulated in the combination treatment group relative to control and IR samples (**Figure 4A-B**). Pathway analysis of up- and down-regulated DEGs using the Hallmark.MSigDB gene set (**Figure 4C, Supplementary Table 6-8**) for combination (Combo, QTP + 4Gy) vs control conditions showed downregulation of G2M checkpoint and E2F targets which is expected for radiation effects on cell cycle and DNA replication pathways, as well as mitotic spindle, Myc targets v1, and spermatogenesis. Among pathways shown to be upregulated in the combination group were cholesterol homeostasis, mTORC1 signaling, TNFα signaling via NF*κβ*, fatty acid metabolism, P53, Hypoxia, complement, androgen response, inflammatory response, and unfolded protein response. Upregulation of these pathways suggest cellular adaptation to combined treatment for cell survival which includes immune response and metabolism regulation.

**Figure 4.**
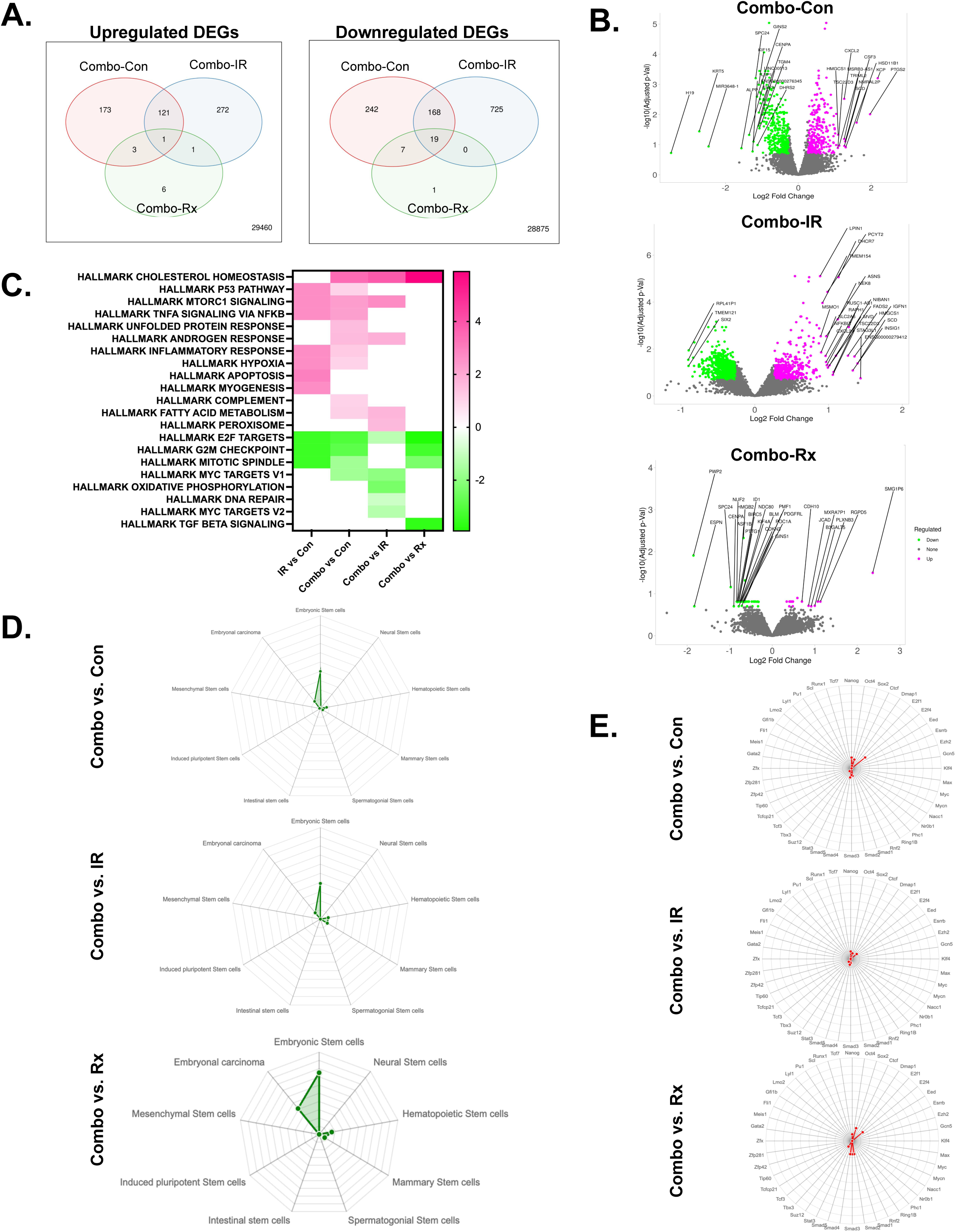
*Combined QTP and radiation treatment upregulates the cholesterol homeostasis pathway and reduces “stemness” in MPM.* RNA-Seq analysis of MSTO-211H MICs 48 hours after treatment with QTP and/or radiation, n=3/treatment group. **A.)** Venn diagrams of DEGs in MSTO-211H cells treated with radiation (IR, 4 Gy), quetiapine (QTP, Rx) or combined QTP+IR (Combo). **B**.) Volcano diagrams of differentially expressed genes (DEGs) in or Combo treated MICs compared with untreated control, IR-treated, or Rx-treated cells with top 25 genes labeled. **C.)** Enrichment analysis using the Hallmark.MSigDB gene set and DEGs. **D.)** Radar charts showing the significance and percentage overlap of the bulkRNAseq gene lists identified via iDEP with Stem Cell datasets in StemChecker and **E.)** transcription factor targets.

Performing a similar pathway analysis for combination treatment versus IR groups, we identified two of the same downregulated pathways as those listed above, with the addition of oxidative phosphorylation, fatty acid metabolism, adipogenesis, MYC targets v2, and DNA repair. Cholesterol homeostasis, mTORC1 signaling, androgen response, fatty acid metabolism, UV Response DN, Peroxisome, and bile acid metabolism were found to be upregulated in the combo group relative to IR. In a pathway analysis for the DEGs from Combo versus Rx (QTP), we found downregulation of G2M checkpoint, E2F Targets, Mitotic Spindle, and TGF*β* signaling. Cholesterol homeostasis was upregulated. As with combination QTP and radiation treatment in GB^14^, we observe similar upregulation of genes involved in the biosynthesis and homeostasis of cholesterol while cell cycle/DNA replication pathways are downregulated following IR.

To confirm whether combined QTP and radiation treatment affects stemness in our MICs, we entered identified downregulated genes into StemChecker ^24^. Most of the queried genes for combo versus control, IR, and Rx groups are expressed in embryonic stem cells and embryonal carcinoma, indicating stem cell signatures are downregulated by combination treatment in MSTO-211H MICs (**Figure 4D**). We also assessed overlap between our gene list and transcription factor (TFs) targets (**Figure 4E**). For combo versus control, TF targets for E2F, SOX2, and NANOG were significantly downregulated. E2F targets were also significantly downregulated in the combo versus IR gene set. No significance was found for combo versus Rx alone. **Supplementary Table 9-10** presents the significance of gene enrichment within composite gene sets associated with various stem cell types and TFs, specifically among the imputed genes identified.

### Single cell RNA sequencing

To evaluate the effects of combination treatment in more detail, we performed single cell RNA sequencing (scRNA-seq) on MSTO-211H MICs treated with 0 Gy (Control, 3,184) or 4 Gy (IR, 2,428), 10 μM QTP (QTP, 3,806), or 10 μM QTP + 4 Gy (QTP_IR, 7,006, **Figure 5**). We confirmed that our cells expressed mesothelial and mesothelioma markers using UMAP visualization plots (**Supplemental Figures 2-3**). Lovain clustering at 0.05 resolution identified 15 unique clusters, some of which cells can be assigned to the different treatment arms of the experiment. We observed an overlap between QTP and QTP+IR cells and—to a lesser extent— IR-treated, QTP-treated, and QTP+IR-treated cells (**Figure 5B**). Furthermore, endothelial- and epithelial-like populations in control, fibroblasts in IR, and tissue stem cell as well as smooth muscle cell-like populations in QTP and QTP_IR groups were annotated (**Figure 5B**). Mesenchymal stem cell (MSC) and neuron-like cells are present in an overlap of all treatment conditions (**Figure 5B**). A shift in cell composition to smooth muscle-like and MSCs, a potential precursor to smooth muscle cells, from endothelial- and epithelial-like cells are observed in QTP and combo treated cells (**Figure 5C-D, Supplementary Tables 11-12**).

**Figure 5.**
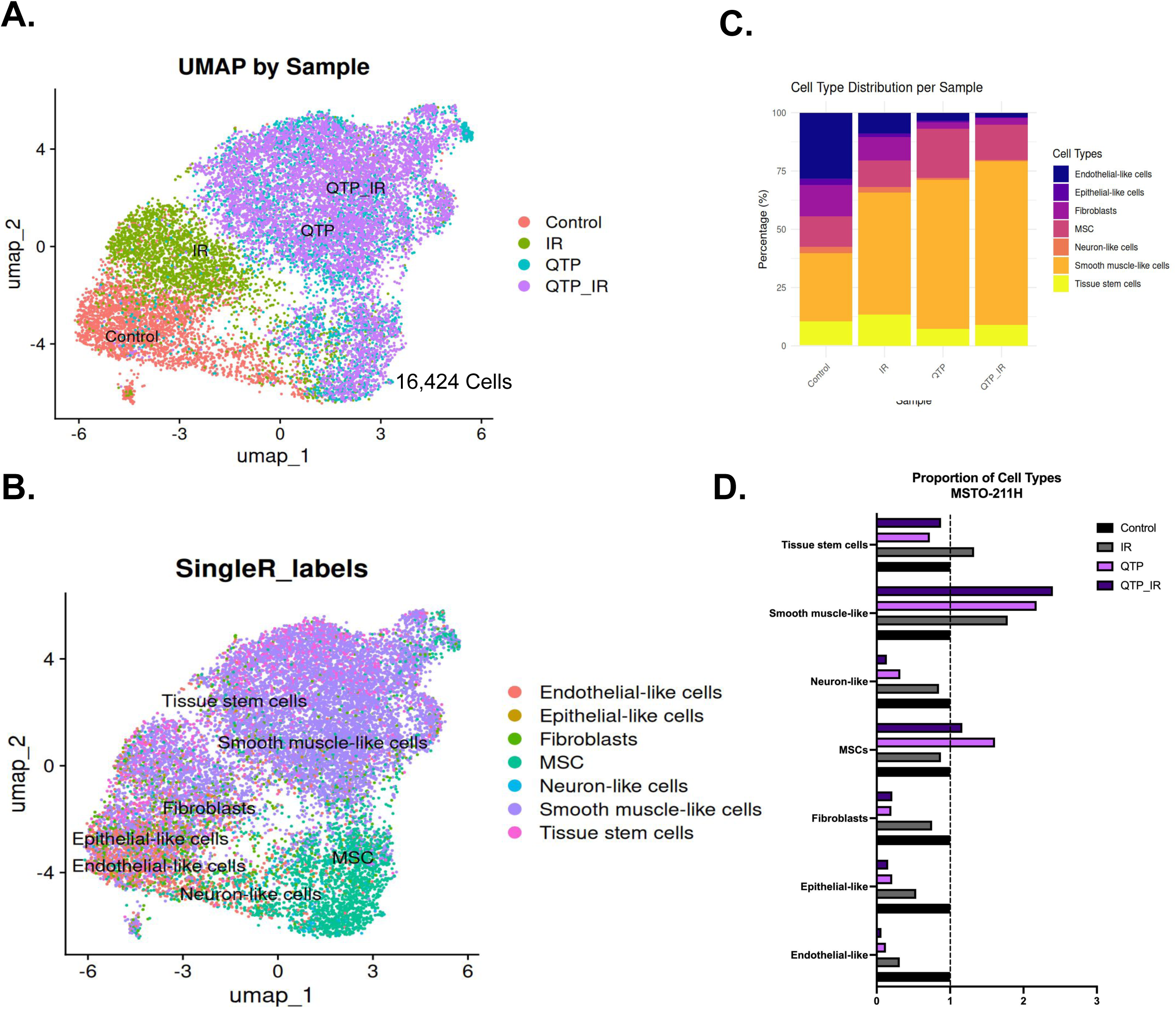
*scRNAseq of treated MSTO-211H MICs 48h post-IR.* **A**.) Uniform Manifold Approximation and Projection (UMAP) plots of clusters identified that could be attributed to the four different treatment groups and **B**.) annotated cell types. **C.)** Stacked columns show percentage (%) distribution of the different cell types identified for each treatment condition. **D.)** Proportion of annotated cell types relative to control.

Using *SenePy* ^25^, we wanted to determine whether QTP+IR increases cellular senescence in our scRNAseq data set (**Figure 6**). We observed significantly higher senescence scores in QTP and QTP+IR groups compared to control and IR groups (**Figure 6B-C**) and expression of senescence markers (**Figure 6D**). Statistics for the distribution of senescence scores can be found in **Supplementary Table 13**. Taken together, our data suggest that QTP and combined QTP and radiation treatment drives cells into cell cycle arrest, potentially inhibiting proliferation of MICs.

**Figure 6.**
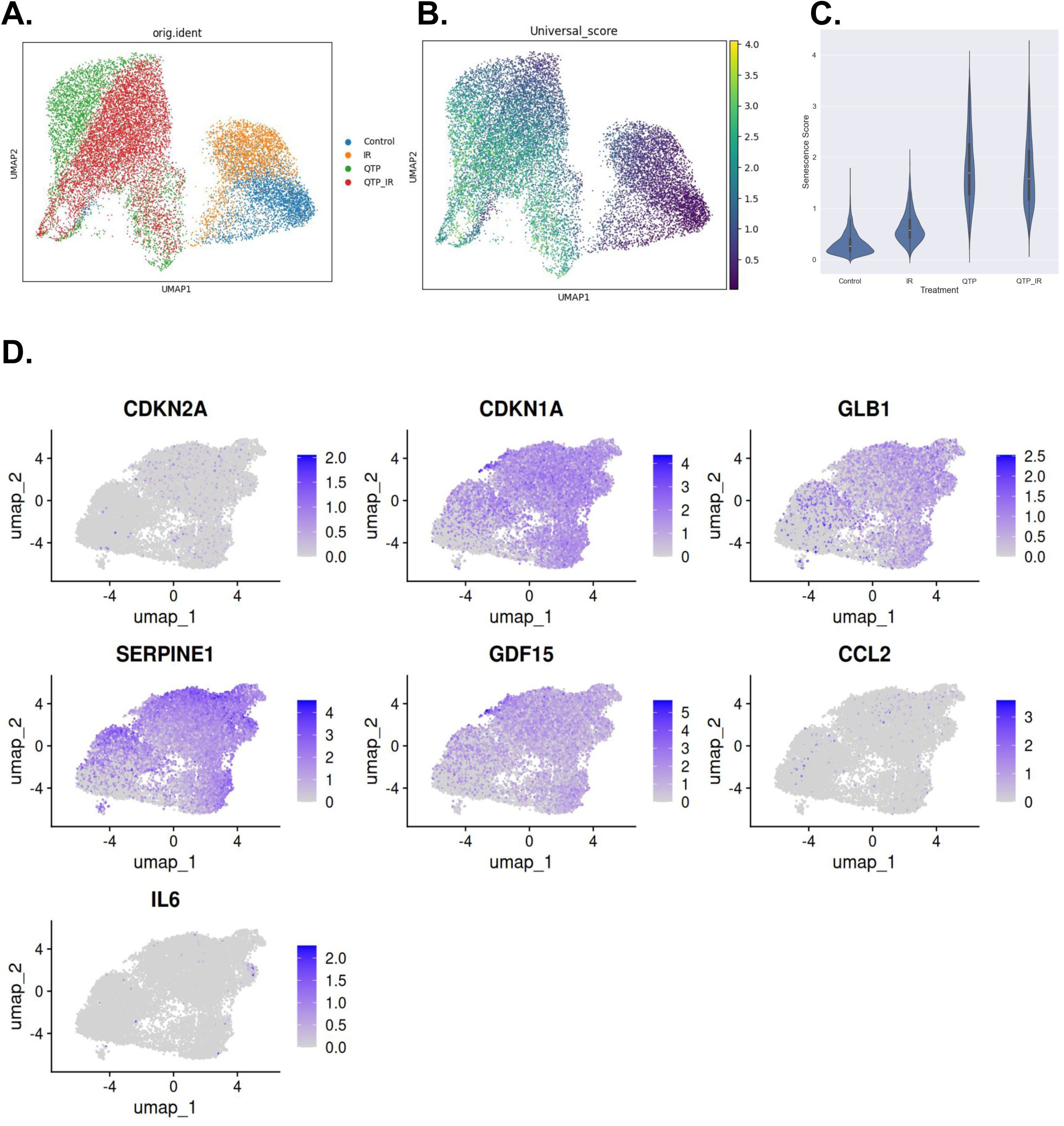
*Cellular senescence in QTP-treated MSTO-211H MICs 48h post-IR.* A.) UMAP plots of clusters identified that could be attributed to the four different treatment groups and **B.)** show the universal score for senescence. **C.)** Violin plot for senescence score separated by treatment group. **D.)** UMAP of senescence marker genes.

## DISCUSSION

In this study, we repurposed the dopamine receptor antagonist, QTP, and tested its combined effects with radiation on MPM. We first show MPM cells express several receptor targets of QTP including DRD2/3 receptors. Combined radiation with QTP treatment reduced self-renewal capacity and stem-cell frequency, but QTP did not radiosensitize MICs. Mechanistically, both QTP alone and in combination with IR altered cellular composition of MPM and induced cellular senescence in MSTO-211H MICs. Importantly, the combination treatment improved survival in a mouse model of MPM.

MPM is a highly heterogenous carcinoma, both histologically and molecularly ^26^. In this study, we leverage both bulkRNA and scRNA sequencing techniques to characterize the cellular response to combined QTP and radiation treatment in mesothelioma spheres, enriched for MICs ^5^. BulkRNAseq analysis reveal an upregulation of stress response pathways and downregulation of cell cycle and proliferation pathways which are typically all upregulated in cultured mesothelioma cells (**Figure 4B, Supplemental Tables 6-8**) ^27^. This is further supported by the downregulation of key TF target genes, including E2F, which regulates cell cycle entry and proliferation in cancer and cancer stem cells ^28, 29^, as well as SOX2 and NANOG which maintain stem-like cell states (Figure 4E) ^30, 31^. Overall, the downregulation of cell cycle and proliferation pathways was in line with the observed increase in senescence associated gene expression profiles observed in our scRNAseq data set.

Cellular senescence is a stress response that can be induced by radiation ^32, 33^ and we show here that the addition of QTP significantly enhances this effect in MICs 48hr post-treatment (**Figure 6**). These findings align with previous reports that QTP promotes cell cycle exit and drives differentiation in primary oligodendrocyte cultures ^34^. While long-term treatment-induced senescence can eventually lead to pro-tumorigenic effects ^32^, our *in vivo* data suggest otherwise (**Figure 3C**). The significantly greater overall survival in QTP+IR-treated animals but not in those receiving QTP alone, indicates that the combined treatment may be effectively controlling tumor growth. Potential mechanisms underlying our observed effects include a true irreversible form of senescence that does not re-enter the cell cycle or alterations in the tumor microenvironment (TME) via senescence-associated secretory phenotype (SASP) factors ^33, 35^ alongside a QTP+IR-induced reduction of a “stem-like” population. Alternatively, pro-tumorigenic effects may be delayed, as the animals ultimately succumb to the tumor ^33^.

ScRNAseq further allowed us to identify distinct cellular subpopulations across treatment groups. Compared to the control sample, QTP and QTP+IR treatment led to a decrease in endothelial-like, epithelial-like, neuron-like, and fibroblast-like populations, while increasing smooth muscle-like cells (**Figure 5C-D**). While studies have demonstrated MPM cells can transdifferentiate into fibroblast-like cells *in vitro* ^36^, we found that QTP and combination treatment reduced fibroblast-like cell populations. Instead, we observed a treatment-induced increase in smooth muscle-like cells which suggest MICs may be transdifferentiating into vascular smooth muscle ^37^. This increase in smooth muscle-like cell populations was most pronounced in the combination treatment group (**Figure 5C-D**). Additionally, the QTP-treated sample showed an increase in MSCs, suggesting a possible mesothelial-to-mesenchymal transition which contributes to tumor progression, therapy resistance, and metastasis and a response to inflammation ^38^ ^39^. This shift in our biphasic MSTO-211H MICs suggests a depletion of the epithelioid cell types and/or a transition towards a more sarcomatoid mesothelioma-like fate from the epithelioid mesothelioma disease state that is less prominent with the addition of RT ^36^.

Tumor-initiating cells or cancer stem cells often are drivers of tumor recurrence and resistance ^35^. In combined QTP+IR treated MICs, we observed a loss of “stemness”, as indicated by significantly reduced SFC (**Figure 1B**), stem cell frequency (**Figure 1C**), expression of stem cell markers (**Figure 4D),** and TF gene targets **(Figure 4E)**. This is further reflected in our scRNAseq data where the combination treatment group had the least distribution and proportion of stem cell populations (**Figure 5C-D**), which may be attributed to a higher proportion of cells undergoing cellular senescence. The observed QTP+IR-induced reduction in “stemness,” coupled with significant survival benefits *in vivo*, indicates that this combined treatment strategy may suppress tumor recurrence and progression, shifting cell populations toward a more differentiated, less aggressive tumor state which would result in better long-term treatment outcomes.

Some limitations of our study include the absence of a well-defined MPM reference dataset for annotating our scRNAseq analysis and the use of the commercially available biphasic cell line MSTO-211H rather than patient-derived models or cell lines from other histological subtypes. Additionally, we sequenced treated MICs grown in suspension without FBS, whereas previous studies primarily focused on adherent cells ^27, 36^. Given the heterogeneity of MPM, future studies should explore the effects of combined QTP+IR treatment across different MPM subtypes (epithelioid and sarcomatoid) and molecular profiles, including both *in vitro* and *in vivo* patient-derived models, and in a mouse model with an intact immune system. Further investigations should also assess the long-term stability of treatment-induced senescence, examine the role of SASP factors, explore the potential integration of senolytic therapies to eliminate senescent cells, and determine whether the observed loss of “stemness” is permanent or transient.

### Conclusion

As radiation therapy techniques continue to advance, integrating QTP with existing treatment modalities could represent a promising approach to improving patient outcomes in MPM ^40, 41^. Further preclinical and clinical studies are warranted to evaluate the translational potential of this combination therapy and its role as a novel MPM treatment strategy.

## Author Contributions

FP and MS conceived the study. FP, LH, and AC contributed to experimental design. FP and AC acquired funding. AC performed the majority of the experiments for this study. EA, LA, TL, KB, and SS performed some experiments. AC and FP analyzed the data. AC and FP drafted the manuscript. All authors contributed to the critical revision of this manuscript.

## Acknowledgements

This work utilized computational and storage services associated with the Hoffman2 Cluster which is operated by UCLA Office of Advanced Research Computing’s Research Technology Group. Preparation of cells for scRNAseq were completed in the UCLA Broad Stem Cell Research Center Sequencing Core. Further we would like to thank Matthew Wong for data management and Carlos Calderon for laboratory assistance.

## Data Availability Statement

All data are available from the corresponding author upon reasonable request.

