## Supplementary Figure 1 for "Combined quetiapine and radiation therapy approach to treat mesothelioma-initiating cells and increase survival in a mouse model of mesothelioma"

### Supplemental Figure 1

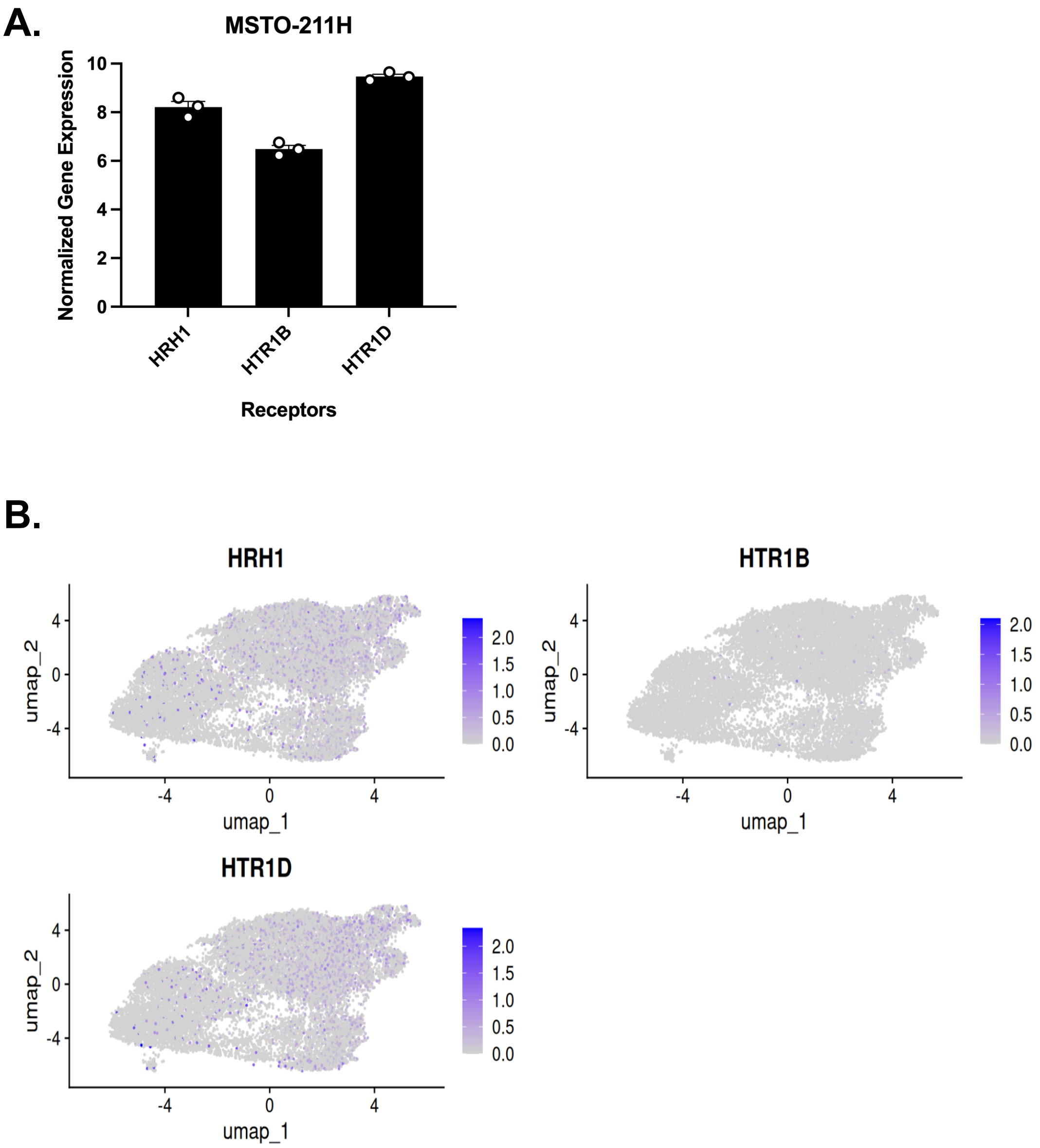


**Supplemental Figure 1.** *QTP receptor expression in MSTO-211H MICs.* **A.)** Normalized baseline gene expression of QTP target receptors in MSTO-211H from bulkRNAseq dataset, n=3 biologically independent repeats +/- SEM. **B.)** UMAP plots of receptor genes from scRNAseq dataset. HRH1 = Histamine 1 Receptor; HTR1B/HTR1D = Serotonin Receptor Subunits 1B/1D.

#### Supplemental Figure 2

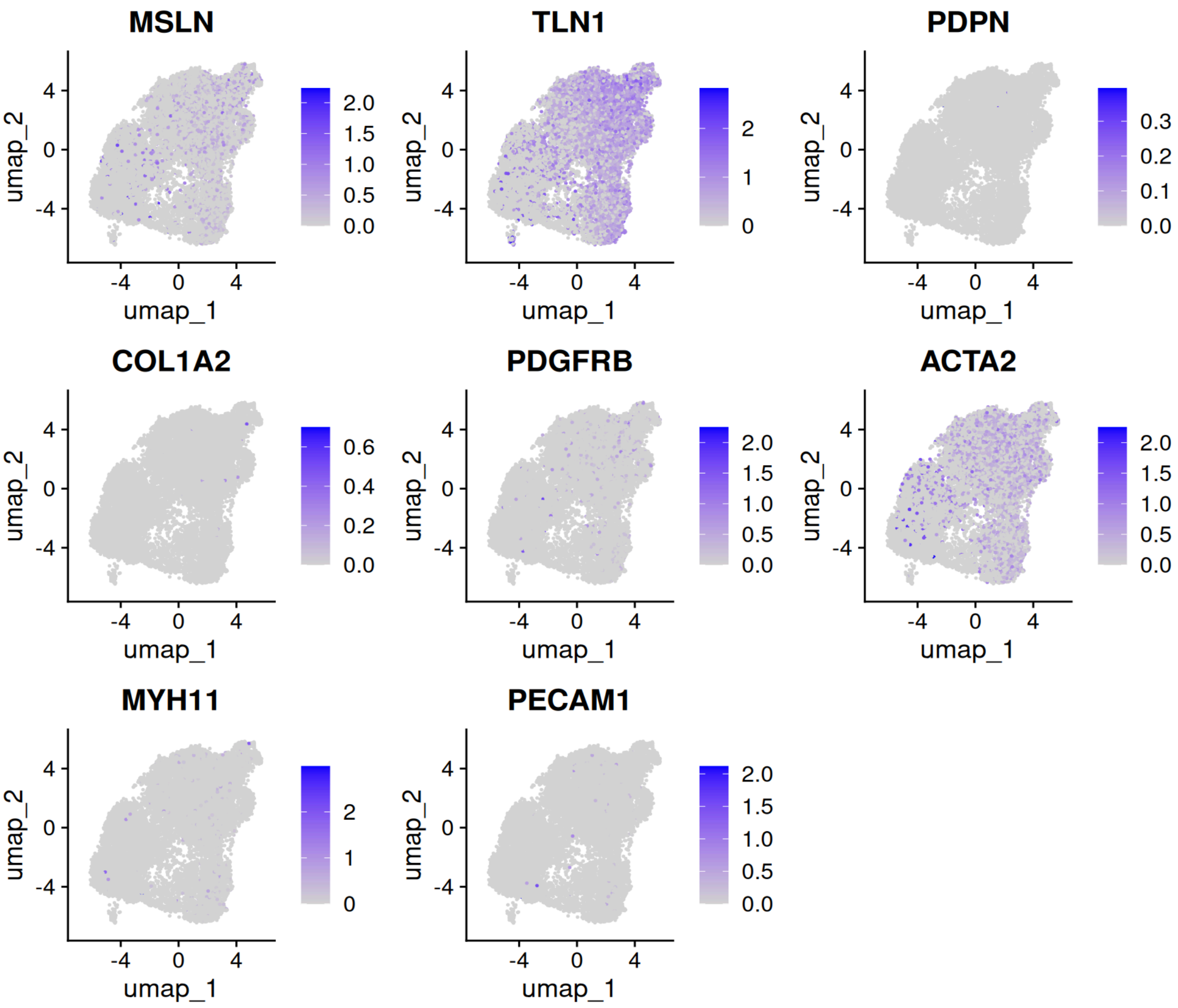


**Supplemental Figure 2.** *UMAP visualization of mesothelial gene marker expression in MSTO-211H MICs.*

#### Supplemental Figure 3

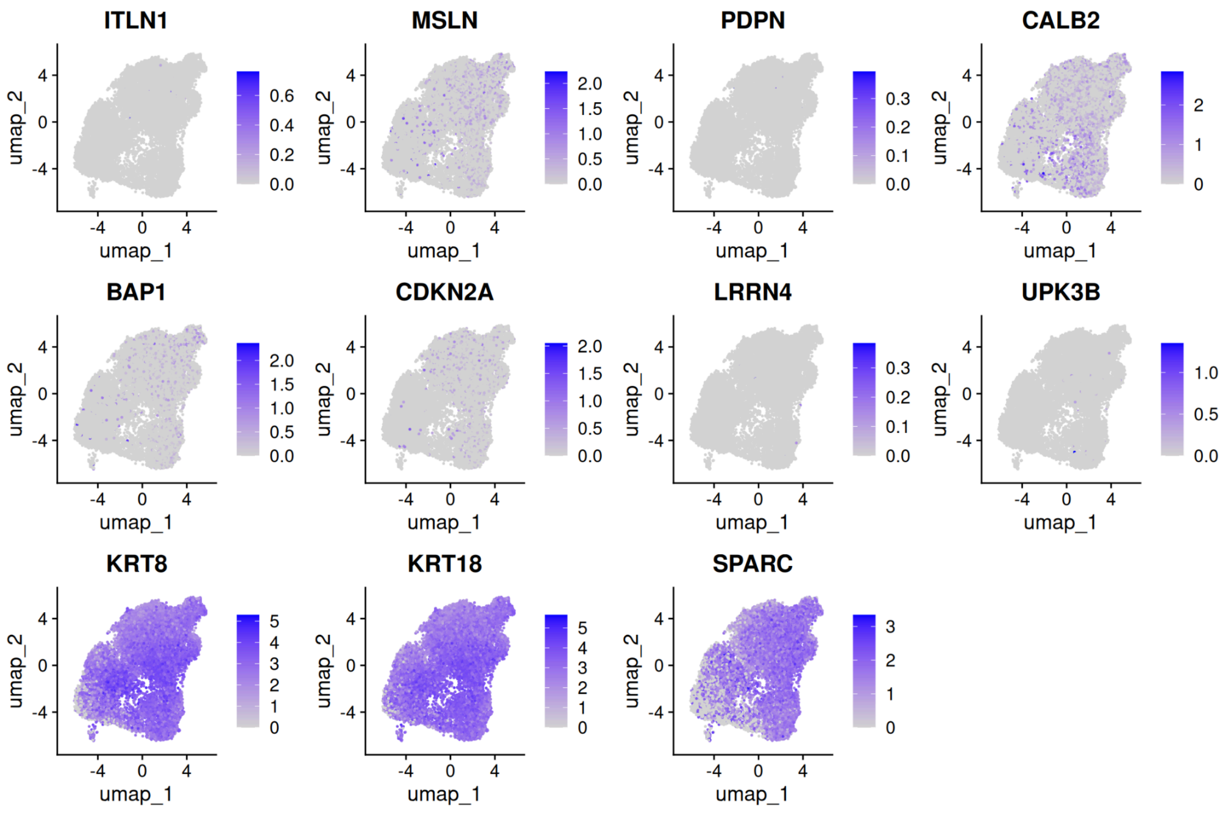


**Supplemental Figure 3.** *UMAP visualization of mesothelioma gene marker expression in MSTO-211H MICs.*

**Supplementary Table 1.** PCR primer sequences (Integrated DNA Technologies) used for RT-PCR experiments.

| Species | Gene Name | Primer sequence (5'-3') |
|---|---|---|
| Human | DRD2 | Forward: TGTACAATACGCGCTACAGCTCCA<br>Reverse: ATGCACTCGTTCTGGTCTGCGTTA |
| Human | DRD3 | Forward: GTGGTGTCCTTCTACCTGCC<br>Reverse: GAGAGAGGGTTTGTTGGGGG |
| Human | PPIA | Forward: ATGCTGGACCCAACACAAAT<br>Reverse: TCTTTCACTTTGCCAAACACC |

**Supplementary Table 2:** Two-way ANOVA analysis results of multiple comparisons for data represented in Figure 1B.

| **MSTO-211H** | | | |
|---|---|---|---|
| **Dunnett's multiple comparisons test** | **Mean Diff** | **95% CI of diff.** | **Adjusted P-value** |
| 0 Gy DMSO vs. 0 Gy QTP (10 μM) | 35.55 | -5.062 to 76.16 | 0.0898 |
| 0 Gy DMSO vs. 4 Gy DMSO | 64.92 | 24.31 to 105.5 | 0.0028** |
| 0 Gy DMSO vs. 4 Gy QTP (10 μM) | 83.29 | 42.68 to 123.9 | 0.0004*** |
| **H2052** | | | |
| 0 Gy DMSO vs. 0 Gy QTP (10 μM) | 66.00 | 34.77 to 97.23 | 0.0003*** |
| 0 Gy DMSO vs. 4 Gy DMSO | 42.75 | 11.52 to 73.98 | 0.0084** |
| 0 Gy DMSO vs. 4 Gy QTP (10 μM) | 78.56 | 47.33 to 109.8 | <0.0001**** |
| **H2452** | | | |
| 0 Gy DMSO vs. 0 Gy QTP (10 μM) | 9.375 | -54.03 to 72.78 | 0.9587 |
| 0 Gy DMSO vs. 4 Gy DMSO | 13.28 | -50.12 to 76.68 | 0.9897 |
| 0 Gy DMSO vs. 4 Gy QTP (10 μM) | 64.93 | 1.527 to 128.3 | 0.0446* |

**Supplementary Table 3:** Confidence Intervals for 1/(stem cell frequency)

| MSTO-211H | | | |
|---|---|---|---|
| **Group** | **Lower** | **Estimate** | **Upper** |
| DMSO_0Gy | 5.8 | 4.69 | 3.82 |
| DMSO_4Gy | 18.3 | 14.59 | 11.68 |
| 10$\mu$M QTP_0Gy | 12.7 | 10.19 | 8.2 |
| 10$\mu$MQTP_4Gy | 34.1 | 27.22 | 21.76 |
| **H2052** | | | |
| **Group** | **Lower** | **Estimate** | **Upper** |
| DMSO_0Gy | 17.9 | 13.9 | 10.8 |
| DMSO_4Gy | 36.6 | 28.3 | 21.8 |
| 10$\mu$M QTP_0Gy | 61.5 | 47.5 | 36.7 |
| 10$\mu$MQTP_4Gy | 143.1 | 110.3 | 85.1 |
| **H2452** | | | |
| **Group** | **Lower** | **Estimate** | **Upper** |
| DMSO_0Gy | 23.3 | 18.6 | 14.9 |
| DMSO_4Gy | 34.6 | 27.6 | 22.0 |
| 10$\mu$M QTP_0Gy | 50.2 | 40.0 | 31.9 |
| 10$\mu$MQTP_4Gy | 125.9 | 100.2 | 79.7 |

**Supplementary Table 4.** Overall test for differences in stem cell frequencies between any of the groups.

| Cell Line | Chisq | DF | P-value |
|---|---|---|---|
| **MSTO-211H** | 142 | 3 | 1.52e-30**** |
| **H2052** | 147 | 3 | 1.35e-31**** |
| **H2452** | 123 | 3 | 1.4e-26**** |

**Supplementary Table 5.** Comparison of survival curves.

| Log Rank test for trend | Chisq | df | P-value |
|---|---|---|---|
| | 8.801 | 3 | 0.0030** |
| **Mantel-Cox Test** | **Chisq** | **df** | **P-value** |
| 0 Gy + Saline vs. 10 Gy + 30 mg/kg QTP | 7.806 | 1 | 0.0052** |
| 10 Gy + Saline vs. 10 Gy + 30 mg/kg QTP | 7.074 | 1 | 0.0078** |
| 0 Gy + 30 mg/kg QTP vs. 10 Gy + 30 mg/kg QTP | 7.589 | 1 | 0.0059** |

**Supplementary Table 6**: Pathway analysis of combo (QTP + 4Gy) versus control DEGs in mesothelioma-initiating cells.

| **Upregulated** | Adj.P-value | Hallmark Signaling Pathways | Fold |
|---|---|---|---|
| | 7.28e-18 | Cholesterol Homeostasis | 10.8 |
| | 7.48e-12 | MTORC1 Signaling | 4.9 |
| | 3.36e-11 | TNFa Signaling via NFKB | 4.7 |
| | 2.15e-02 | Unfolded Protein Response | 3 |
| | 2.16e-02 | Androgen Response | 3 |
| | 2.16e-02 | Inflammatory Response | 2.3 |
| | 8.15e-02 | Hypoxia | 2 |
| | 8.15e-02 | Complement | 2 |
| | 8.15e-02 | P53 Pathway | 2 |
| | 8.91e-02 | Fatty Acid Metabolism | 2.1 |
| **Downregulated** | Adj.P-value | Hallmark Signaling Pathways | Fold |
| | 3.09e-59 | E2F Targets | 8.8 |
| | 5.98e-43 | G2M Checkpoint | 7.4 |
| | 1.8e-7 | Mitotic Spindle | 3.2 |
| | 1.8-7 | MYC Targets V1 | 3.2 |
| | 7.52e-2 | Spermatogenesis | 2.1 |

**Supplementary Table 7**. Pathway analysis of combo versus IR (4 Gy) DEGs in mesothelioma-initiating cells.

| **Upregulated** | Adj. P-value | Hallmark Signaling Pathways | FDR |
|---|---|---|---|
| | 1.15e-22 | Cholesterol Homeostasis | 14 |
| | 5.44e-17 | MTORC1 Signaling | 6.4 |
| | 1.8e-3 | Fatty Acid Metabolism | 3.3 |
| | 8.04e-3 | Androgen Response | 3.6 |
| | 1.45e-2 | Bile Acid Metabolism | 3.2 |
| | 2.02e-2 | UV Response DN | 2.8 |
| | 2.43e-2 | Peroxisome | 3.1 |
| **Downregulated** | Adj. P-value | Hallmark Signaling Pathways | FDR |
| | 5.54e-35 | Oxidative Phosphorylation | 5.2 |
| | 3.36e-20 | MYC Targets V1 | 4 |
| | 4.69e-5 | E2F Targets | 2.3 |
| | 4.44e-4 | Fatty Acid Metabolism | 2.3 |
| | 7.55e-3 | Adipogenesis | 1.9 |
| | 3.49e-2 | DNA Repair | 1.9 |
| | 4.32e-2 | MYC Targets V2 | 2.5 |

**Supplementary Table 8**. Pathway analysis of combo versus Rx (QTP) DEGs in mesothelioma-initiating cells.

| **Upregulated** | Adj. P-value | Hallmark Signaling Pathways | FDR |
|---|---|---|---|
| | 1.69e-2 | Cholesterol Homeostasis | 59.2 |
| **Downregulated** | Adj. P-value | Hallmark Signaling Pathways | FDR |
| | 4.4e-9 | E2F Targets | 15.2 |
| | 5.5e-5 | G2M Checkpoint | 10.1 |
| | 3.55e-2 | Mitotic Spindle | 12.5 |
| | 4.71e-2 | TGFB Signaling | 5.1 |

**Supplementary Table 9**. Enrichment of genes included in composite gene sets for the different stem cell types among the imputed downregulated genes. Significance (p-value) is calculated by the hypergeometric test. Adjusted p-value is calculated by Bonferroni correction.

| Combo vs. Con | | | | |
|---|---|---|---|---|
| **Cell Type** | **Set Size** | **Overlapping genes** | **P-value** | **Adjusted P-value** |
| Embryonic Stem Cells | 2933 | 103 | 1e-16 | 9e-16**** |
| Neural Stem Cells | 168 | 4 | 0.192 | 1 |
| Hematopoietic Stem Cells | 967 | 20 | 0.039 | 0.35 |
| Mammary Stem Cells | 305 | 8 | 0.056 | 0.501 |
| Induced Pluripotent Stem Cells | 80 | 2 | 0.294 | 1 |
| Mesenchymal Stem Cells | 114 | 4 | 0.069 | 0.619 |
| Embryonal Carcinoma | 635 | 28 | 4.046e-8 | 3.641e-7**** |
| **Combo vs. IR** | | | | |
| **Cell Type** | **Set Size** | **Overlapping genes** | **P-value** | **Adjusted P-value** |
| Embryonic Stem Cells | 2977 | 237 | 1e-16 | 9e-16**** |
| Neural Stem Cells | 168 | 5 | 0.678 | 1 |
| Hematopoietic Stem Cells | 962 | 60 | 3.973e-6 | 3.575e-5**** |
| Mammary Stem Cells | 301 | 63 | 1e-16 | 9e-16**** |
| Induced Pluripotent Stem Cells | 80 | 1 | 0.937 | 1 |
| Mesenchymal Stem Cells | 114 | 5 | 0.345 | 1 |
| Embryonal Carcinoma | 643 | 60 | 1.199e-12 | 1.079e-11**** |
| **Combo vs. Rx** | | | | |
| **Cell Type** | **Set Size** | **Overlapping genes** | **P-value** | **Adjusted P-value** |
| Embryonic Stem Cells | 3021 | 9 | 9.37e-6 | 8.437e-5**** |
| Hematopoietic Stem Cells | 968 | 2 | 0.123 | 1 |
| Mammary Stem Cells | 306 | 1 | 0.178 | 1 |
| Embryonal Carcinoma | 650 | 5 | 3.078e-5 | 2.77e-4**** |

**Supplementary Table 10.** Enrichment of genes included in composite transcription factor (TF) gene sets for the different TF gene targets among the imputed downregulated genes. Significance (p-value) is calculated by the hypergeometric test. Adjusted p-value is calculated by Bonferroni correction.

| Combo vs. Con | | | | |
|---|---|---|---|---|
| **Set Name** | **Set Size** | **Overlapping genes** | **P-value** | **Adjusted P-value** |
| E2F_Boyer1 | 1038 | 51 | 5.631e-16 | 5.631e-15**** |
| SOX2_Boyer1 | 1156 | 29 | 9.692e-4 | 0.01** |
| NANOG_Boyer1 | 1511 | 34 | 0.002 | 0.023* |
| SMAD2_Brown | 128 | 6 | 0.008 | 0.079 |
| SMAD3_Brown | 128 | 6 | 0.008 | 0.079 |
| OCT4_Boyer1 | 566 | 10 | 0.236 | 1 |
| SMAD3_Kim | 1266 | 16 | 0.645 | 1 |
| SMAD4_Kim | 2373 | 30 | 0.679 | 1 |
| SMAD2_Kim | 1619 | 18 | 0.837 | 1 |
| SUZ12_Lee | 1509 | 11 | 0.993 | 1 |
| **Combo vs. IR** | | | | |
| **Set Name** | **Set Size** | **Overlapping genes** | **P-value** | **Adjusted P-value** |
| E2F_Boyer1 | 1038 | 59 | 7.201e-5 | 7.201e-4**** |
| NANOG_Boyer1 | 63 | 63 | 0.051 | 0.513 |
| SOX2_Boyer1 | 47 | 47 | 0.113 | 1 |
| SMAD2_Brown | 5 | 5 | 0.439 | 1 |
| SMAD3_Brown | 5 | 5 | 0.439 | 1 |
| OCT4_Boyer1 | 13 | 13 | 0.95 | 1 |
| SMAD3_Kim | 29 | 29 | 0.993 | 1 |
| SUZ12_Lee | 29 | 29 | 1 | 1 |
| SMAD2_Kim | 31 | 31 | 1 | 1 |

| | | | | |
|---|---|---|---|---|
| SMAD4_Kim | 48 | 48 | 1 | 1 |
| **Combo vs. Rx** | | | | |
| **Set Name** | **Set Size** | **Overlapping genes** | **P-value** | **Adjusted P-value** |
| SMAD2_Brown | 128 | 1 | 0.078 | 0.784 |
| SMAD3_Brown | 128 | 1 | 0.078 | 0.784 |
| E2F_Boyer1 | 1038 | 2 | 0.138 | 1 |
| SOX2_Boyer1 | 1156 | 2 | 0.165 | 1 |
| SMAD4_Kim | 2373 | 2 | 0.456 | 1 |
| SMAD3_KIm | 1266 | 1 | 0.565 | 1 |
| SUZ12_Lee | 1509 | 1 | 0.632 | 1 |
| NANOG_Boyer1 | 1511 | 1 | 0.632 | 1 |
| SMAD2_Kim | 1619 | 1 | 0.659 | 1 |

**Supplementary Table 11.** Percentage composition by treatment group for SingleR-labeled cells in our dataset.

| SingleR_labeled cells | Control | IR | QTP | QTP_IR |
|---|---|---|---|---|
| Endothelial-like | 28.2882 | 8.8759 | 3.5438 | 1.8497 |
| Epithelial-like | 2.7690 | 1.4931 | 0.5861 | 0.4335 |
| Fibroblasts | 13.3417 | 10.0788 | 2.6645 | 2.8323 |
| MSCs | 13.0899 | 11.4475 | 21.1031 | 15.2601 |
| Neuron-like | 2.7375 | 2.3226 | 0.8792 | 0.3757 |
| Smooth muscle-like | 29.2951 | 52.3019 | 63.8689 | 70.3323 |
| Tissue stem cells | 10.1636 | 13.4798 | 7.3541 | 8.9161 |

**Supplementary Table 12**. Cell composition (count) for SingleR-labeled cells in our dataset.

| SingleR_labeled cells | Control | IR | QTP | QTP_IR |
|---|---|---|---|---|
| Endothelial-like | 899 | 214 | 133 | 128 |
| Epithelial-like | 88 | 36 | 22 | 30 |
| Fibroblasts | 424 | 243 | 100 | 196 |
| MSCs | 416 | 276 | 792 | 1056 |
| Neuron-like | 87 | 56 | 33 | 26 |
| Smooth muscle-like | 931 | 1261 | 2397 | 4867 |
| Tissue stem cells | 323 | 325 | 276 | 617 |

**Supplementary Table 13:** One-sided Mann Whitney U test for distribution of senescence scores.

| **Comparison** | **Mann-Whitney U Statistic** | **P-value** |
|---|---|---|
| Con vs IR | 1301819.5 | 0.0**** |
| Con vs QTP | 98871.5 | 0.0**** |
| Con vs QTP+IR | 169745 | 0.0**** |
| IR vs QTP+IR | 895313 | 0.0**** |
| QTP vs QTP+IR | 13293856 | 1.63e-11**** |
